# Uncovering structural features that control substrate specificity in a Lactobacillus chlorogenic acid esterase

**DOI:** 10.1101/2023.05.30.542953

**Authors:** Kellie K. Omori, Tracie L. S. Okumura, Nathaniel B. Carl, Brianna T. Dinn, Destiny Ly, Kylie N. Sacapano, Allie Tajii, Cedric P. Owens

**Author notes:** Corresponding author: Cedric Owens, Schmid College of Science and Technology, Department of Chemistry and Biochemistry, Chapman University, 1 University Dr., Orange, CA 92866.

## Abstract

The structural determinants of chlorogenic acid esterase (CE) substrate specificity are poorly understood. Here, we establish how a *Lactobacillus helveticus* CE selects for its substrate, chlorogenic acid (CGA). We determine that a Lys residue in an extended loop over the active site imparts substrate specificity by hydrogen bonding to CGA. Mutation of the Lys residue abolishes CGA specificity. Comparison with other bacterial CEs reveals that the extended loop is not conserved. However, the hydrogen bonding functionality to CGA is preserved thanks to other residues. Structural comparison with ferulic acid esterases (FAEs), a related enzyme class, shows that CEs feature a more restricted active site, reflecting the fact that they hydrolyze smaller substrates compared to FAEs.

## Introduction

Ferulic acid esterases (FAEs, also known as cinnamoyl esterases, EC 3.1.1.73) are a class of enzymes that are active against a broad range of hydroxycinnamic acid esters [1-3]. FAEs act on substrates such as coumaric acid esters [3], ferulic acid polysaccharides [3], and chlorogenic acid (CGA) [4, 5], each differing in terms of their substitutions around the benzene ring and the size and polarity of the group that is further from the ester carbonyl [1]. FAEs are important biotechnologically and in the context of health and nutrition. Hydroxycinnamic acids are used in the cosmetics industry [6] and are being investigated for their anti-cancer and anti-diabetic potential [7]. As phenolic compounds, hydroxycinnamic acid derivatives are proposed to be beneficial to human health because of their antioxidant properties and their ability to modulate the growth of beneficial bacterial in the human gut [7-9]. FAE treatment of hydroxycinnamic acid-rich food is therefore being investigated as a way of altering the phenolic composition in food to increase antioxidant uptake in the human diet and to improve the gut microbiome [9, 10].

FAEs are found in both fungi and bacteria. All known FAEs are α/β hydrolases, a superfamily of enzymes that share a common α/β fold and possess a conserved Ser-His-Asp catalytic triad. Since FAEs cleave diverse hydroxycinnamic acid derivatives, there is considerable structural variety between FAEs. Currently, FAEs are empirically classified based on their substrate preference [2, 11]. Such classification, however, does not provide information on the specific structural features that determine why a certain FAE preferably hydrolyzes one of many similar looking substrates. Moreover, to date, most structure-function studies on FAEs have focused on understanding how the enzyme selects substrates based on the substitutions around the hydroxycinnamic acid’s aromatic ring [2, 12]. In contrast, there is much less information on the mechanism of by which FAEs recognize substrates based on the properties of the group further from the ester carbonyl (the group that extends out of the active site pocket). Such knowledge, however, is critical for understanding FAE function as enzyme-substrate interactions outside of the core active site pocket have a significant impact on substrate specificity [13-15]. Notably, a study on an *Aspergillus niger* FAE (AnFaeA) revealed that the catalytic efficiency for the natural substrate arabinose ferulate was 1600-fold higher than for a methyl ferulate model substrate. [15]

Chlorogenic acid esterases (also referred to as chlorogenate esterases, CEs, EC 3.1.1.42) are closely related to FAEs and referred to interchangeably in the literature as their substrate range overlaps [5, 16]. CEs hydrolyze CGA, a water-soluble ester of caffeic and quinic acid. CEs also hydrolyze other substrates such as ethyl- and methyl ferulic acid [17]. CGA is a beneficial dietary antioxidant but sometimes undesired in food due to its taste and because it can cause some foods to discolor [18, 19]. Thus, CEs are being investigated for their ability to improve the sensory properties of foods such as coffee and sunflower flour [18, 20]. This work focuses on understanding the structural factors that govern CGA binding in CEs and to compare how substrate selection differs in CEs and FAEs.

The only bacterial CE that has been both structurally and functionally characterized is a CE from *Lactobacillus johnsonii* (Lj0536) [5, 21]. Lj0536 has a slight preference for CGA over the smaller substrate ethyl ferulic acid. The specificity constant, *k*_cat_/*K*_m_, for CGA hydrolysis is 1.4- fold higher for CGA [5]. Like many α/β hydrolases, Lj0536 contains an insertion or lid domain.

The insertion domain sits above an elongated active site cleft and has an α-ββ-ββ-α architecture [21]. Mutational analysis in the Lj0536 insertion domain showed that mutations that abolish H-bonding between the enzyme and the caffeic acid moiety of CGA affected both *k*_cat_ and *K*_m_ [21]. Notably, a Gln_145_ to Ala mutation near the top of the active site cleft decreased both CGA and ethyl ferulic acid hydrolysis. The mutation’s effect was greater for CGA, suggesting this residue may be involved in CGA binding. However, the reason for the differential effect of the Gln_145_ to Ala mutation was not further explored.

Our group recently characterized a putative CE from *Lactobacillus helveticus* (LhCE). LhCE was very active against CGA and was used to break down CGA in sunflower flour to prevent sunflower flour cookie discoloration [18], however, its substrate specificity was never unambiguously determined [16, 22]. The primary sequence of LhCE is similar to that of Lj0536 (**Figure S1**). However, the LhCE α/β insertion domain is slightly different from the one in Lj0536 since it contains a short insertion (Asn_163_-Lys_168_) in a region that is predicted to be a loop directly above the active site cleft [22]. The extended loop in the insertion domain is rare for bacterial CEs. An NCBI BLAST search found only two other CEs that feature the longer loop, one in *L. gallinarum*, and another *L. crispatus* (**Figure S1**). The large size of the loop raises the question of how it will affect CGA binding and hydrolysis in LhCGA. On the one hand, the larger size may restrict access for CGA. On the other, it may stabilize binding of CGA by offering additional H-bonding opportunities to the quinic acid moiety in the active site. Either way, a comparison of LhCE structure and activity with Lj0536 will provide useful information on the mechanism of substrate recognition in CEs.

This study presents the structure of LhCE and determines that LhCE is more catalytically active against CGA compared to smaller hydroxycinnamic acid ester substrates. Docking studies indicate that the Lys on the extended loop stabilizes CGA binding through H-bonding. Mutation of the loop Lys residues weakens CGA binding and lowers catalytic specificity towards CGA. Structural comparisons with *L. johnsonii* CEs support the conclusion that H-bonding between quinic acid and residues at the entrance to the active site are essential for CGA binding. Further analyses indicate that CEs distinguish themselves from related FAEs by featuring more restricted active sites that enhance CGA binding. Overall, this study reveals key structural features that enable CGA binding to CEs and highlights how studying closely related CEs can determine the origins of their substrate selectivity.

## Methods

### Materials

Materials were purchased from Sigma-Aldrich, Fisher Scientific and VWR. All reagents were ACS grade or equivalent. Primers were purchased from IDT.

### Cloning and mutagenesis

The K164A mutant was made by site directed mutagenesis using wild type LhCE as template with the Q5 Site-Directed Mutagenesis Kit from NEB using following primers:

Forward: GGTGGGCAACGCGCTGGGCATGAAAGTGGG

Reverse: AGCGGCACCACATCCGGA

Pet28a plasmids containing the esterase gene were cloned into BL21(DE3) cells, grown overnight in LB broth containing 50 μg/mL kanamycin. The plasmid was then purified using a miniprep kit (Thermo). Plasmids were sent for sequencing (Genewiz) to confirm the presence of the mutation.

### Protein expression, purification, and physical characterization

Protein expression and purification for WT and K164A LhCE were carried out as described in Lo Verde *et al*. [22]. The gel filtration, circular dichroism and thermal stability measurements were also carried out as described in Lo Verde *et al*.[22].

### Protein crystallization

Crystallization of WT and K164A LhCE were both performed using the hanging drop vapor diffusion method with a well volume of 1 mL and a drop size of 2 μL. The drop contained 1 μL of protein and 1 μL well solution. For WT LhCE, the well solution contained 0.1 M citric acid, pH 5.0, 10% PEG 6000, 1.6 M lithium chloride. The protein concentration was 20 mg/mL. For K164A LhCE, the well solution contained 0.1 M citric acid pH 5.0, 7.5% PEG 6000, 0.01 M zinc chloride, and 1.5 M lithium chloride. The protein concentration was 19 mg/mL. For both proteins, crystals formed after approximately two weeks, and crystals were harvested and cryo-protected in a solution containing equal volume well solution and 50% glycerol, and flash frozen in liquid nitrogen.

### Data collection and structure determination

WT LhCE was diffracted at ALS beamline 12-2 and K164A LhCE at ALS beamline 5-02 using a wavelength of 1 LJ. Indexing, scaling, merging, and truncation was done using the Dials pipeline [23] in Xia2. For WT LhCE, two datasets were merged prior to molecular replacement. The structure of WT LhCE was solved by molecular replacement with 3PF8 as the search model. WT LhCE was used as the search model for K164A LhCE. After molecular replacement, an initial structure was generated using Phenix Autobuild [24]. Structural refinement was conducted iteratively using Phenix Refine and Coot [25]. All protein structures were visualized in Pymol [26].

### Docking simulations

Docking models were created using Molsoft ICM. Default settings for ligand docking were used and thoroughness/effort was 3 for all complexes. The models with the lowest (best) ICM scores were selected for further analysis.

### Kinetic characterization of LhCE

The kinetic properties of WT and K164A LhCE were determined using Michaelis-Menten assays. Hydrolysis of CGA, ethyl caffeic acid, and ethyl ferulic acid were all carried out at 21 °C using an Agilent Technologies Cary 60 UV-Vis spectrophotometer in a buffered solution of 50 mM HEPES, pH 8. A stock solution of CGA was prepared by dissolving CGA in 50 mM HEPES pH 8.0. Stock solutions of ethyl caffeate and ethyl ferulic acid were made by dissolving each substrate in 50 mM HEPES pH 8.0, 5% ethanol. The concentration of each solution was calculated using the Lambert-Beer law using their respective extinction coefficients (**Figure S2**). For all experiments, the enzyme concentration was 10 nM and the reaction volume was 0.8 mL.

All reactions were monitored for one minute at 0.05-minute intervals from 250 nm to 400 nm. When the substrate concentrations were under 0.10 mM, the pathlength was 1 cm. For concentrations above 0.10 mM, the pathlength was 0.2 cm. The equations used for calculating product formation from the difference in molar absorptivity are listed in the supporting information. All kinetics data represents the average of at least four true independent replicates with at least three different batches of enzyme. Enzymatic velocities were calculating using a linear fit of the initial rates, and enzyme kinetics parameters calculated by nonlinear curve fitting. All kinetic data analysis was conducted in GraphPad Prism 9.

## Results and Discussion

### Structure of WT LhCE and comparison with Lj0536

The structure of WT LhCE was solved by single-crystal x-ray diffraction to a resolution of 2.2 Å with R_free_/R_work_ values of 0.21 and 0.18, respectively (**Figure 1A**). The crystallography statistics are summarized in **Table 1**. The asymmetric unit of LhCE contains three chains, with two chains forming a dimer. The third chain forms a dimer with its counterpart in a neighboring asymmetric unit. The LhCE dimer interface is large, approximately 2910 Å^2^, consistent with the fact that the protein is dimeric in solution [22]. The overall structure of LhCE is typical of the α/β hydrolase fold, containing at its core eight β stands connected by α helices. The α/β insertion domain starts at Ala_132_ and ends at Leu_180_ and is located above the active site cleft (**Figure 1A** and **B**).

**Table 1.**
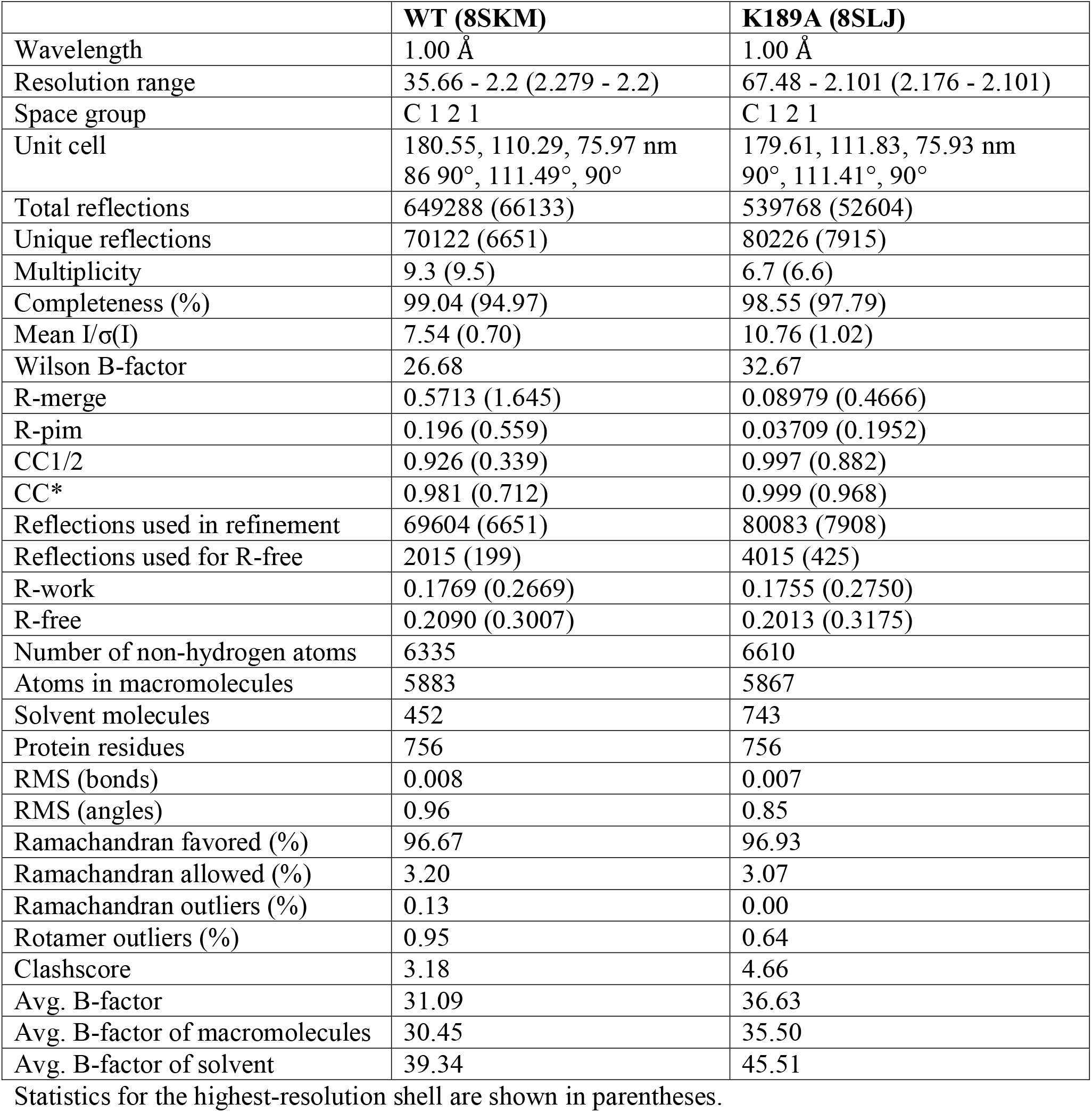
Data collection and refinement statistics.

**Figure 1.**
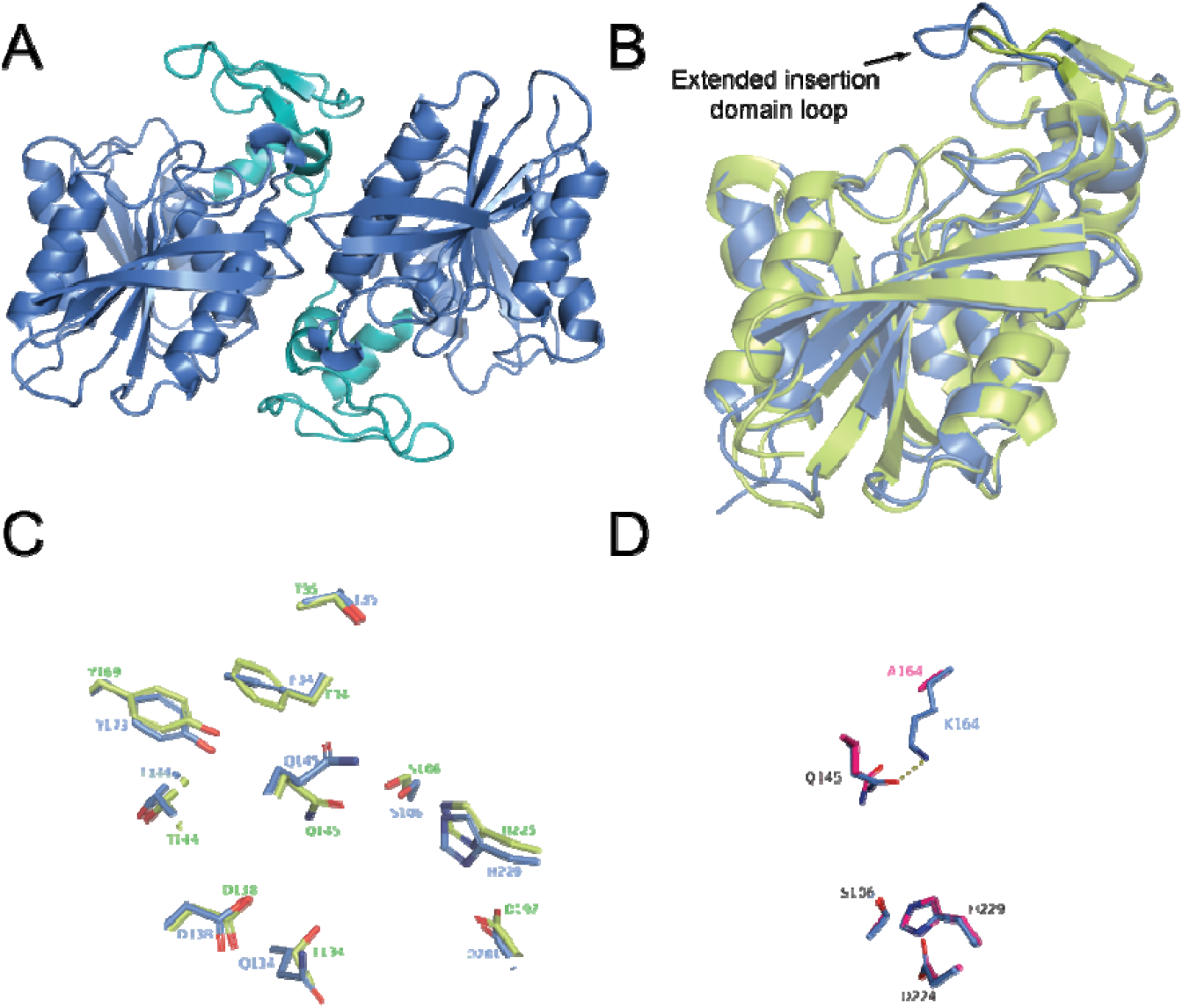
(A) Structure of the LhCE dimer. The insertion domain is highlighted in teal. (B) Structural comparison of the protomers of LhCE (blue) and Lj0536 (PDB ID: 3PF8, green). (C) Close up of the active sites of LhCE (blue) and Lj0536 (green), highlighting the close structural similarity. The catalytic triad is composed of Ser_106_, His_225_ and Asp_197_ (LhCE numbering). (D) Comparison of WT LhCE (blue) and K164A LhCE (pink) that demonstrates that except for residue K164, the two structures are superimposable. The H-bond between Lys_164_ and Gln_145_ i also shown in this image.

LhCE and Lj0536 are highly similar with and RMSD of less than 1 □ over the aligned Cα (**Figure 1B**). The active site residues are highly conserved (**Figure 1C**), as are the active site pocket volumes, as calculated in Molsoft (283 Å^3^ for LhCE and 291 Å^3^ for Lj0536). The largest structural difference between LhCE and Lj0536 is the extension of the active site loop in LhCE above the active site, as shown in **Figure 1B**. The extended loop protrudes about 6 □ further over the active site in LhCE compared Lj0536. In LhCE, the extended loop contains a lysine (Lys_164_). It forms a 2.8 □ H-bond with residue LhCE Gln_145_ which is located on a loop below LhCE Lys_164_ (**Figure 1D**). Lj0536 also has a lysine residue in the loop above the active site, however, due to the loop’s smaller size, the lysine residue sits outside of the active site cleft in Lj0536. Based on the LhCE structure, is seems likely that Lys_164_ will be able to H-bond with the quinic acid moiety in CGA, thereby increasing CGA binding specificity.

### Structure of K164A LhCE

A Lys_164_ to Ala mutant was generated to study the role of the extended loop in LhCE. Protein expression and purification for the mutant was identical to WT LhCE and its physical propertie were the same as the wild type enzyme (**Figure S3**). The structure of K164A LhCE was solved to 2.1 Å and R_free_/R_work_ values of 0.20 and 0.18, respectively (**Table 1**). It is identical to WT LhCE with an RMSD of 0.165 Å per protomer (**Figure 1D and S4**). The active site is also identical, the only difference being the absence of Lys_164_ in the loop of the mutant (**Figure 1D**). There are no significant differences in the sidechain orientation of loop and active site residues in the K164A- and WT LhCE structures except for a single residue, Gln_145_. Since this residue no longer can H-bond with Lys_164_, it moves horizontally approximately 2 □ compared to the wild-type structure (**Figure 1D**).

### Docking CGA to LhCE suggests Lys_164_ stabilizes CGA binding

To date, there are no experimental structures of CGA bound to a CE, however, a Lj0536 structure with ethyl ferulic acid has been reported [21]. Our attempts to obtain substrate-bound structures by soaking LhCE crystals with CGA yielded only apo-protein. Thus, we turned to docking studies to understand how substrate binding is influenced by the extended loop. Docking was performed using Molsoft ICM [27] with WT LhCE and K164A LhCE. To start, we tested the accuracy of our docking process by generating ethyl ferulic acid bound to Lj0536, which recapitulated the crystal structure [21] (**Figure S5**).

CGA docked deeply into LhCE the active site cleft and the ester is oriented so that it can be attacked the by the hydroxide of the active site Ser (**Figure 2A**). The caffeic acid moiety of CGA binds to the active site in a similar fashion as ferulic acid binds in the Lj0536 crystal structure. Numerous H-bonds to the caffeic acid moiety and ester stabilize CGA binding. Two noteworthy short H-bonds are between the hydroxyl groups of the catechol ring and Asp_161_ and Tyr_196_. In addition, Phe_57_ may form a 3.9 □ π-stacking interaction with CGA catechol ring. **Table 2** show a list of key interactions involved in CGA binding.

**Table 2.** Amino acids involved in H-bonding or ionic interactions with CGA in docking models.

|  | No. of H- bonds to CGA | Amino acid residues involved |
| --- | --- | --- |
| WT LhCE | 13 | D138, Y173, S106, H229, Q145, F34, T35, A36, R230, K164 |
| K187A LhCE | 10 | D138, Y173, S106, H229, Q145, F34, T35, A36 |
| Lj0536 | 13 | D138, Y169, H225, Q145, H105, F34, T35, A36, G33, Q107 |

**Figure 2.**
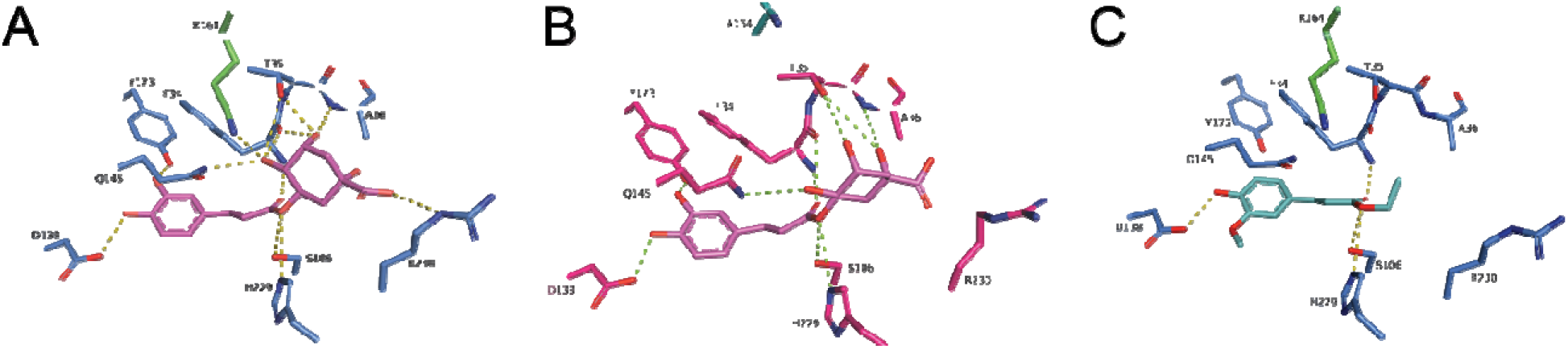
Docking models for (A) CGA binding to LhCE, (B) CGA binding to K164A LhCE, (C) ethyl ferulic acid binding to LhCE. Key residues are shown as sticks and important H-bonds within 4 □ of the substrate are shown as dashed lines.

As predicted, Lys_164_ H-bonds to the quinic acid moiety of CGA (**Figure 2A**). Gln_145_, Thr_35_ and the main chain of several residues, including Phe_34_, are also involved in a hydrogen bonding network with quinic acid. In contrast, the K164A mutant lacks the H-bond to quinic acid from residue 164 (**Figure 2B**). There is a slight rotation of the quinic acid in the docked K164A LhCE structure, leading to a loss of H-bonds with Thr_35_ and the main chain of Phe_34_ in the mutant docked complex (**Figure 2B** and **Table 2**).

The structures of ethyl ferulic acid (**Figure 2C**) and ethyl caffeic acid (**Figure S6**) bound LhCE are very similar to the caffeic acid end of the CGA structure as the interactions around the aromatic ring are essentially the same for all three substrates. As expected, neither substrate interacts with Lys_164_ (**Figure S6**), further supporting the hypothesis that the role of Lys_164_ is to specifically stabilize CGA binding.

### Enzyme kinetics data demonstrating Lys_164_ enables CGA specificity

To determine the importance of Lys_164_ on CGA binding and turnover, the Michaelis-Menten parameters for WT and K164A LhCE were measured and are shown in **Figure 3A** and **Table 3**. For WT LhCE, the kinetic constants are within the range of error of our previously published values [22]. The data shows that the *K*_m_ of K164A LhCE towards CGA is 2.6-fold higher than the *K*_m_ of the WT, strongly supporting the hypothesis that the loop Lys is important for CGA binding. Surprisingly, *V*_max_ of the mutant also increases and is 1.4-fold higher than that of the WT. The difference is statistically significant with a p-value of 0.022. These experiments were the average of seven independent triplicate runs so we are confident that *V*_max_ indeed increases in the K164A mutant. We hypothesize that this unexpected outcome is the result of the H-bond between Lys_164_ and Gln_145_, which may keep the loop in a closed conformation, thus limiting the rate of substrate entry and/or release. Alternately, Lys_164_ may simply represent a steric barrier to substrate release. Although *V*_max_ increased, the mutant is nevertheless the worse enzyme for CGA hydrolysis as the *k*_cat_/*K*_m_ value decreased 1.9-fold in the mutant compared to the wild type. These results indicate that Lys_164_ is important for maximizing CGA hydrolysis efficiency in LhCE.

**Table 3.** Michaelis-Menten parameters for CGA, ethyl caffeic and ethyl ferulic acid hydrolysis for WT and K164A LhCE.

| | $K_m$ (mM) | $V_{max}$<br>($\mu\text{mol min}^{-1}\text{mg}^{-1}$) | $V_{max}$<br>($\text{mM min}^{-1}\text{mg}^{-1}$) | $k_{cat}$<br>($\text{s}^{-1}$) | $k_{cat}/K_m$<br>($\text{mM}^{-1}\text{s}^{-1}$) |
| --- | --- | --- | --- | --- | --- |
| **WT** |  |  |  |  |  |
| CGA | $0.147 \pm 0.041$ | $164.6 \pm 38$ | $205.8 \pm 47.6$ | $82.6 \pm 19.1$ | 561.8 |
| ethyl ferulate | $0.386 \pm 0.046$ | $177.1 \pm 18.4$ | $221.4 \pm 23$ | $88.9 \pm 9.2$ | 230.2 |
| ethyl caffeate | $0.645 \pm 0.112$ | $499.9 \pm 79.9$ | $624.9 \pm 99.8$ | $250.8 \pm 40.1$ | 388.7 |
| **K187A** |  |  |  |  |  |
| CGA | $0.384 \pm 0.111$ | $227 \pm 49.4$ | $283.8 \pm 61.7$ | $113.9 \pm 24.8$ | 296.5 |
| ethyl ferulate | $0.38 \pm 0.077$ | $166.7 \pm 19.8$ | $208.4 \pm 24.8$ | $83.6 \pm 10$ | 219.9 |
| ethyl caffeate | $0.564 \pm 0.163$ | $460.2 \pm 117.1$ | $575.2 \pm 146.3$ | $230.9 \pm 58.7$ | 409.0 |

**Figure 3.**
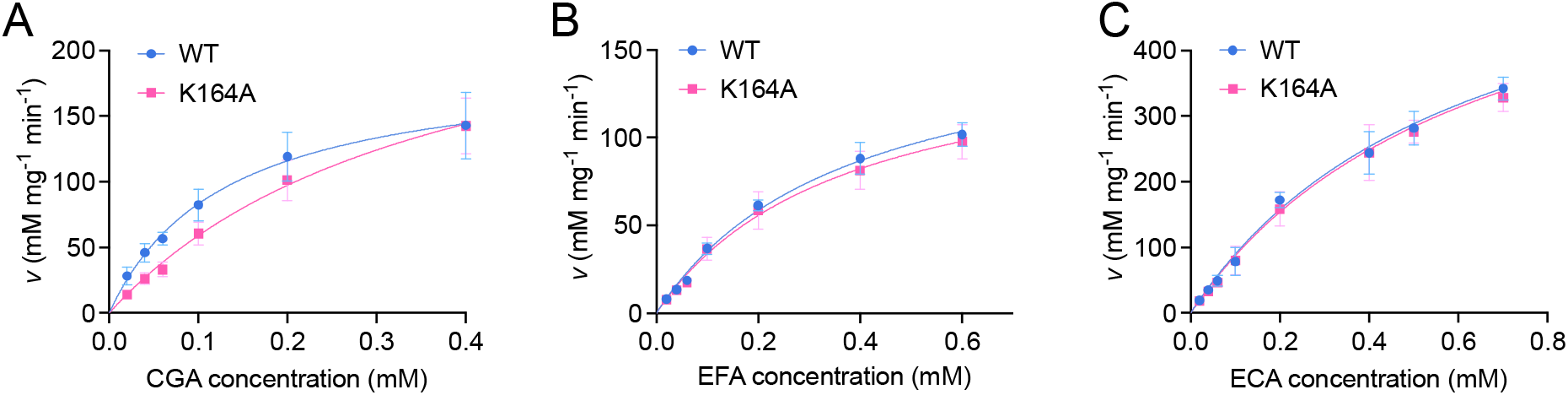
Michaelis-Menten analysis for (A) CGA, (B) ethyl ferulic acid (EFA), and (C) ethyl caffeic acid (EFA) hydrolysis. Each graph represents the average of at least five independent triplicate measurements. The kinetic constants for these experiments are listed in Table 3. We note that the error for the *K*_m_ for ethyl caffeic acid is large since saturation was not reached du to the enzyme’s weak affinity for the substrate.

To tests if the mutation at Lys_164_ affects only CGA catalysis or influences activity in general, we carried out kinetics experiments with the WT and K164A LhCE with ethyl caffeic and ethyl ferulic acid. We predicted that the *V*_max_ and *K*_m_ values would be indistinguishable between WT LhCE and K164A LhCE. The values of *K*_m_ and values of *V*_max_ between WT and K164A LhCE are within error of each other for both ethyl caffeic acid and ethyl ferulic acid, supporting this hypothesis (**Figure 3B and C** and **Table 3**). Furthermore, the respective *K*_m_ values for ethyl ferulic and ethyl caffeic acid are higher than those for CGA, which is consistent with the idea that the role of the extended loop is to enhance CGA binding.

The LhCE *k*_cat_ values indicate that turnover is fastest for ethyl caffeic acid (**Table 3**). We note, however, that the *k*_cat_/*K*_m_ value is nevertheless highest for CGA. A potential reason for the higher *k*_cat_ for ethyl caffeic acid hydrolysis is that product release after ester cleavage is faster for ethanol than for quinic acid due the former’s much smaller size and the fact that ethanol does not H-bond with Lys_164_. The reason for the slower turnover of ethyl ferulic acid compared to ethyl caffeic acid is likely related to the size of the aromatic ring. Ethyl caffeic and ethyl ferulic acid differ by methylation of C3 alcohol of the catechol ring. This causes the width of the ring to be 7.5 □ for ethyl ferulic acid, compared to 5.25 □ for ethyl caffeic acid, meaning ferulic acid release may require greater opening of the active site cleft to allow product release. Taken together, the kinetics data allows us to conclude the following: 1) LhCE is indeed a CGA esterase, as proposed in literature [16, 22]. 2) Enzyme specificity depends on Lys_164_ to strengthen CGA binding. 3) Ethyl caffeic acid hydrolysis features the highest *k*_cat_, suggesting that tighter binding of the larger CGA and ethyl ferulic acid substrates come at the cost of a slower turnover rate.

### Comparison between LhCE and LjCE CGA binding

Having established that Lys_164_ is important for stabilizing CGA binding in LhCE, we were interested in finding CGA-specific H-bonding interactions in Lj0536 (**Figure 4A**). In addition, CGA binding to an Alphafold [28] model of Lj1228 was also tested (**Figure 4B**). Lj1228 is a *L. johnsonii* CE that is known to have a strong CGA preference and is has high sequence homology to Lj1228 (**Figure S1**) [5]. As expected, the docked CGA structures for LhCE, Lj0536 and Lj1228 are very similar to each other with the overall orientation of the substrates being the same. Both the Lj0536 and Lj1228 docking models reveal H-bonds that stabilize the quinic acid moiety of CGA. For Lj1228, the H-bond originates from residue Lys_35_, which is located on a loop at the side of the active site cleft (**Figure 4B**). Its sidechain amine is positioned roughly where the Lys_164_ in the extended loop in LhCE would be located. CGA in Lj0536 is stabilized through H-bonds to quinic acid that originate from both Gln_145_, which is located on a loop above the active site, and His_105_ (**Figure 4A**). Lj0536 His_105_ is located adjacent to the catalytic Ser_106_. The interaction between His_105_ and CGA is reminiscent of the one between His_233_ and arabinofuranosyl ferulic acid in a docking model with a *Lentilactobacillus buchneri* (Lb) FAE [29]. Like His_105_ in Lj0536, His_233_ in LbFAE is located adjacent to the active site Ser and was suggested to aid in holding the substrate in place. It is also worth noting that the lysine residue in the Lj0536 loop (Lys_161_) is not within H-bonding distance of CGA. However, it may still contribute to binding by weakly interacting with the carboxylic acid on quinic acid.

**Figure 4.**
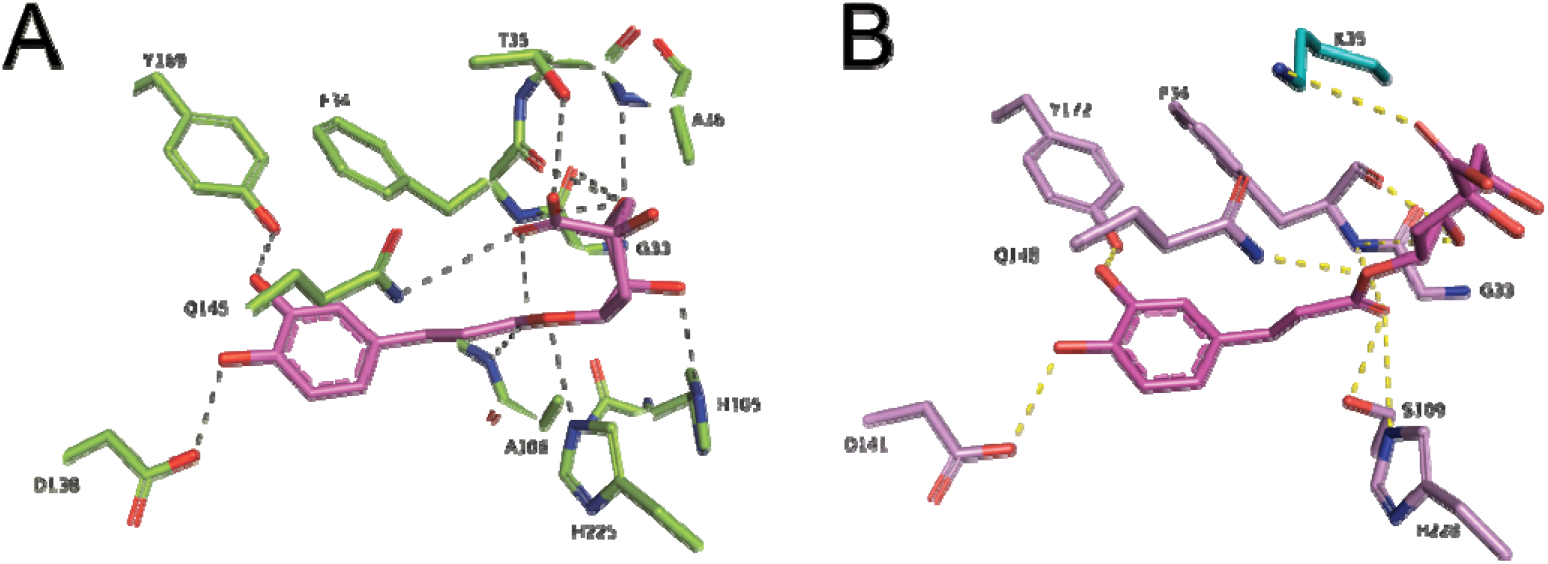
Docking models for (A) CGA binding to Lj0536 (PDB ID 3S2Z) and (B) an Alphafold model of Lj1228. Key residues are shown as sticks and important H-bonds within 4 □ of th substrate are shown as dashed lines.

The stabilizing interactions mentioned above only occur with CGA since Lj1228 Lys_35_ and Lj0536 Gln_145_ and His_105_ do not H-bond to either ethyl caffeic or ethyl ferulic acid in both the docking model and, for Lj0536, in a previously published co-crystal structure [21]. This suggest that these interactions are functionally similar to the ones between LhCE Lys_164_ and CGA. Importantly, discovering that residue Gln_145_ H-bonds to CGA in Lj0536 provides an explanation for the observation that the Lj0536 Q145A mutation had a more detrimental effect on CGA hydrolysis than on ethyl ferulic acid cleavage.

### Structural comparison of bacterial CEs with related FAEs

The data thus far shows that H-bonding at the entrance to the active site cleft is important for imparting CGA binding specificity in CEs. The next question we wished to answer was if there are structural features that set bacterial CEs apart from FAEs. Both CEs and FAEs cleave ethyl ferulic acid and ethyl caffeic acid, suggesting that determining the preferred substrate must occur on the periphery of the respective esterase’s active site pocket.

Surface representations of the CEs, LhCE, Lj0536 and Lj1228 are shown in **Figure 5A** and enlarged in **Figure S7**. They each feature an enclosed binding pocket around the caffeic acid moiety of CGA with quinic acid protruding from the pocket through a collar. In all three cases, the insertion domain and α/β core forms a relatively narrow cleft for the substrate to enter. There are only few structures of bacterial FAEs in the PDB. Four well-characterized FAEs are shown in **Figure 5B**. They all feature a typical α/β hydrolase core. However, the insertion domains for CEs and FAEs are very different. In the case of a *Lactobacillus plantarum* FAE (Lp0796) [30] and *Alistipes shahii* FAE (AsFAE) [31], the insertion domain is α-helical. In Lp0796 it is spatially removed from the α/β hydrolase core, whereas in AsFAE, the domain does not form a closed lid over the active site. In both cases, this leads to the binding pocket being open and easily accessible. In *Dysgonomonas mossii* FAE (DmFAE), there is no insertion domain. Instead, the region above the active site is composed of a separate β-sandwich carbohydrate binding domain, leading to a very large active site [32]. Furthermore, in a *Bacteroides intestinalis* (Bi) FAE, the insertion domain is small, consisting only of an extended, flexible loop [14]. While these four proteins are structurally distinct, they all share a solvent exposed active site pocket in common. All four active sites are much more open than those in CEs. We hypothesize that the open active site allows FAEs to act on their large, carbohydrate-decorated hydroxycinnamic acid substrates, whereas CEs have evolved to act on the smaller substrate CGA.

**Figure 5.**
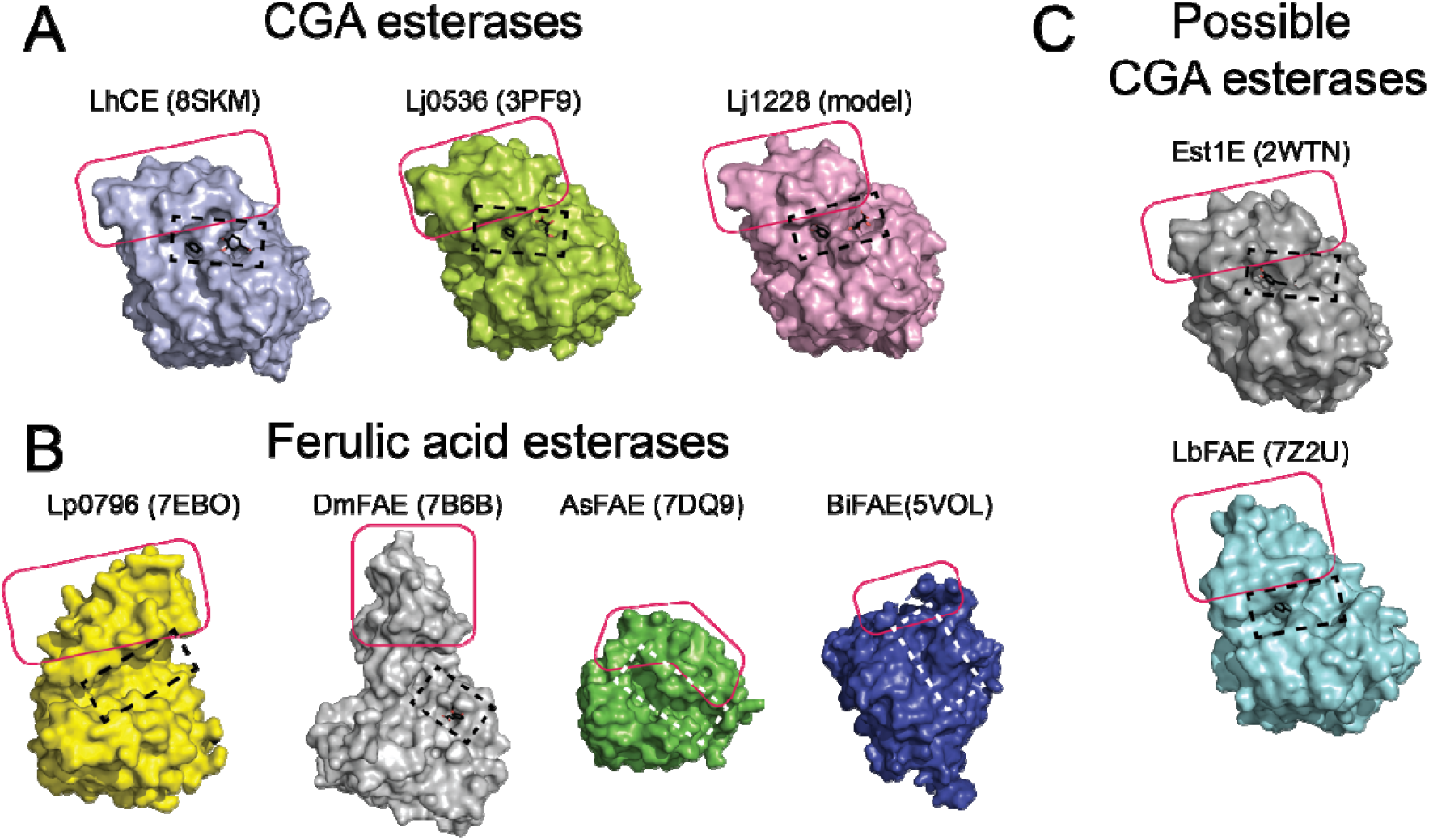
(A) Surface representations of LhCE (8SKM), Lj0536 (3S2Z), and Lj1228 (Alphafold model). (B) Surface representations of the FAEs Lp0796 (7EBO), DmFAE (7B6B), and BiFAE (5VOL) and (C) of Est1E (2WTN) and LbFAE (7Z2U). The CGA ligand in LhCE, Lj1228 and Lj0536 are from our docking studies. The ferulic acid ligands in Est1E, LbFAE and DmFAE were found in the experimental structure. The insertion domain, loop, or carbohydrate binding domain in DmFAE are boxed in red. The active site is highlighted by a dashed box. Note that all proteins shown here except for AsFAE are dimers. Only a single protomer is displayed for clarity. The active site in LbFAE is located near the dimer interface, which may further restrict substrate access.

Finally, this structural comparison raises the question of the true identity of *Butyrivibrio proteoclasticus* Est1E [33] and a *Lentilactobacillus buchneri* FAE (LbFAE) [29], both shown in **Figure 5C**. Although both enzymes are highly homologous to LhCE with RMSD values of ca. 1 □ over the aligned Cα (**Figure S8**), they are currently classified as FAEs. CGA hydrolysis activity was never assayed in either enzyme. However, given their high structural homology to *Lactobacillus* CEs and similar active site pocket structure, it seems likely that Lp0796 and Est1E will have higher activity against CGA compared to smaller hydroxycinnamic acid esters.

## Conclusions

The data presented herein indicate that substrate specificity in CEs is derived from H-bonding between the quinic acid moiety and residues located at the top of the CE’s active site cleft. The importance of the H-bond between enzyme and CGA was demonstrated by mutational analysi in LhCE, which indicated that removing the H-bond lowers the substrate binding affinity, causing a loss in specificity. Importantly, our data for K164A LhCE is consistent with observations from a previously described Q145A Lj0536 mutant [21]. In both enzymes, removing the stabilizing H-bond to the quinic acid moiety of CGA leads to a loss of CGA hydrolysis. The results further indicate that the residues involved in H-bonding with CGA do not have to be conserved since H-bonding involves a lysine residue in LhCE (Lys_164_) and Lj1228 (Lys_35_) and a His residue in Lj0536 (His_105_). Overall, these results support the notion that CEs have adopted similar, but slightly divergent structural strategies to ensure substrate preference for CGA. A limitation of our study is that substrate selection in CEs may not be explained entirely by H-bonding between enzyme and substrate. Fluctuations in the insertion domain as a whole likely alter the size of the opening to the active site cleft, thereby influencing substrate binding, as suggested by the EstE1 crystal structure in complex with ethyl ferulic acid [33]. Our group is currently investigating this possibility through studies that probe the insertion domain’s conformational changes under turnover. Finally, we compared the structures of bacterial CEs and FAEs, discovering that these enzymes, while similar, feature active sites that differ substantially in terms of solvent exposure. This comparison suggests that studying CEs and FAEs with model substrates may not capture the full substrate recognition mechanism which likely depends on more than just the hydroxycinnamic acid moiety.

## Abbreviations

CE: chlorogenic acid esterase
CGA: chlorogenic acid
ECA: ethyl caffeic acid
EFA: ethyl ferulic acid
FAE: ferulic acid esterase

## Author contributions

## Investigation and Editing

Kellie K. Omori, Tracie L. S. Okumura, Nathaniel B. Carl, Brianna T. Dinn, Destiny Ly, Kylie N. Sacapano, Allie Tajii, Cedric P. Owens; **Conceptualization, Supervision, Writing:** Kellie Omori, Cedric P. Owens; **Funding acquisition:** Cedric P. Owens

## Acknowledgments

The authors would like to thank Dr. Marco Bisoffi for use of instrumentation in his labs and Drs. Andy Borovik, Nicholas Chim, and James Nowick for sharing their beamtime with our group. We are grateful to Dr. Lilian W. Senger for fruitful discussion and acknowledge her collaboration on NIFA grant 2020-67018-31261 that supported part of this project.

## Funding

This work was supported by USDA-NIFA Novel Foods and Innovative Manufacturing Technologies grant 2020-67018-31261 to C.P.O., National Science Foundation, Division of Chemistry grant 1905399 to C.P.O, and a Research Corporation Cottrell Scholar Award, and a Postbac Scholar award to C.P.O. K.K.O acknowledges support by the Chapman Center for Undergraduate Excellence.

## Notes

### Competing Interest Statement

The authors have declared no competing interest.

