## Supporting information for "Uncovering structural features that control substrate specificity in a Lactobacillus chlorogenic acid esterase"

^*^ Corresponding author:

Cedric Owens

**Supplementary methods**

Equation for determining hydrolysis products spectrophotometrically based on experimentally determined extinction coefficients:

$$\Delta\left[ product \right]= \frac{A-A_{o}}{\varepsilon_{product}-\varepsilon_{susbtrate}} (Equation 1)$$

Where the products are caffeic acid or ferulic acid, as indicated in the main text.

The extinction coefficients that were used are as follows at λ = 340 nm:

Chlorogenic acid: 13957 (mM cm)^-1^

Ethyl caffeic acid: 8848 (mM cm)^-1^

Ethyl ferulic acid: 8895 (mM cm)^-1^

Caffeic acid: 3590 (mM cm)^-1^

Ferulic acid: 4328 (mM cm)^-1^

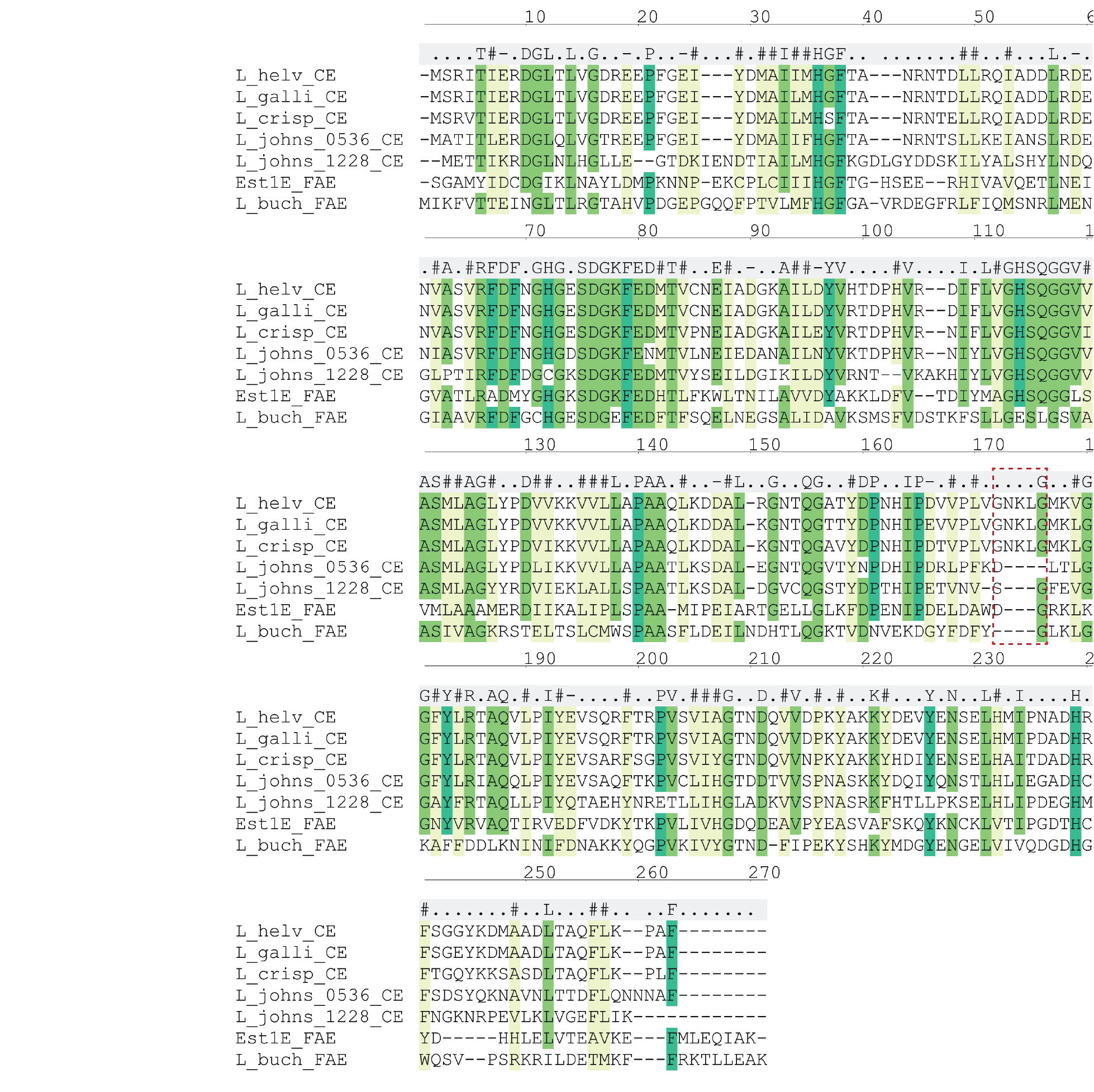

**Figure S1.** Sequence alignment between *L. helveticus* CE and related CEs and FAEs. Consensus residues are shown at the top of the alignment. The extended insertion domain loop is highlighted by the dashed red box and starts at consensus sequence residue 171.

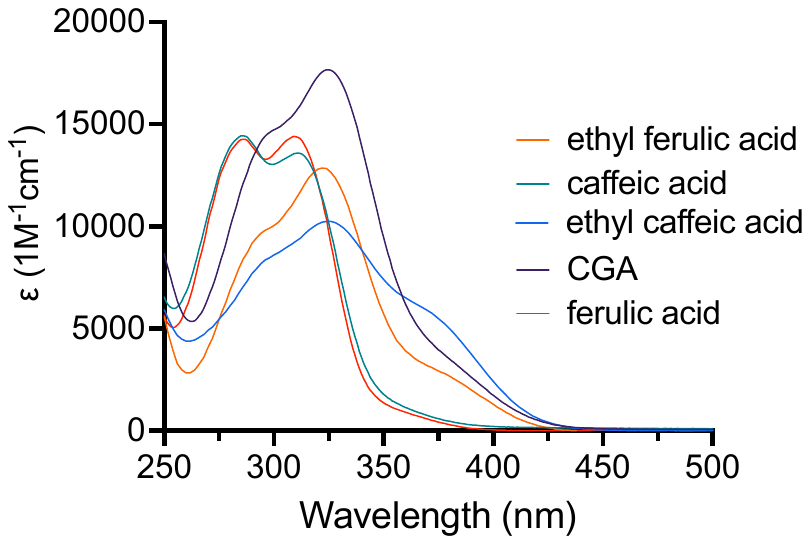

**Figure S2.** Experimentally determined extinction coefficients for the substrates and products used in this study. The average of at least four independent replicates were used to calculate the extinction coefficients.

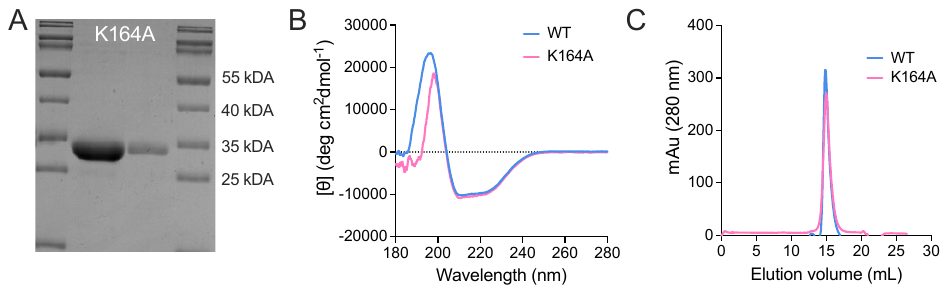

**Figure S3.** (A) SDS PAGE (10%) of K164A LhCE at two different concentrations, demonstrating that the protein is pure. (B) Circular dichroism spectra of WT and K164A LhCE indicating that the secondary structure is identical for both proteins. (C) Gel filtration chromatogram of WT and K164A LhCE demonstrating that their molecular weight is the same. Proteins (2 mg) were run on a S200 10/30 column in a buffered solution of 50 mM HEPES, pH 8 at a flow rate of 0.7 ml/min. The elution volume is consistent with that of a LhCE dimer (60 kDa) based on a calibration curve with molecular weight standards.

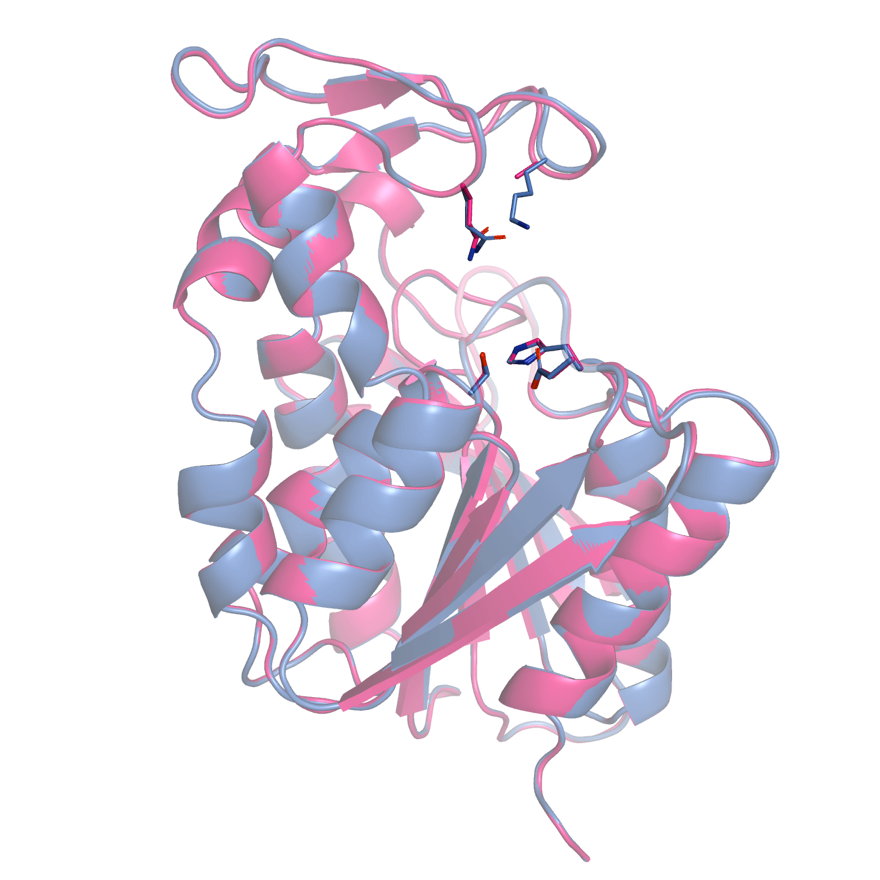

**Figure S4.** Superposition of a protomer of WT and K164A LhCE demonstrating that the structures are nearly identical. WT LhCE is colored blue and K164A LhCE is pink.

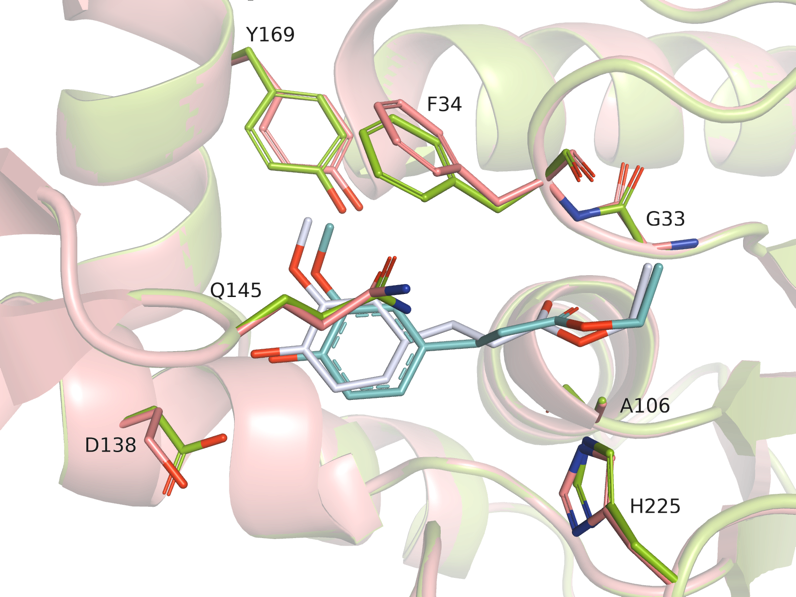

**Figure S5.** Control for Molsoft ICM docking studies. Ethyl ferulic acid was docked into Lj0536 and compared to the experimentally reported co-crystal structure (PDB ID 3QM1). In the docking model, the polypeptide is green and ethyl ferulic acid is teal. In the experimental structure, the polypeptide is pink and the ligand is white. The docked ligand binds to Lj0536 nearly in the same orientation as in the experimental structure.

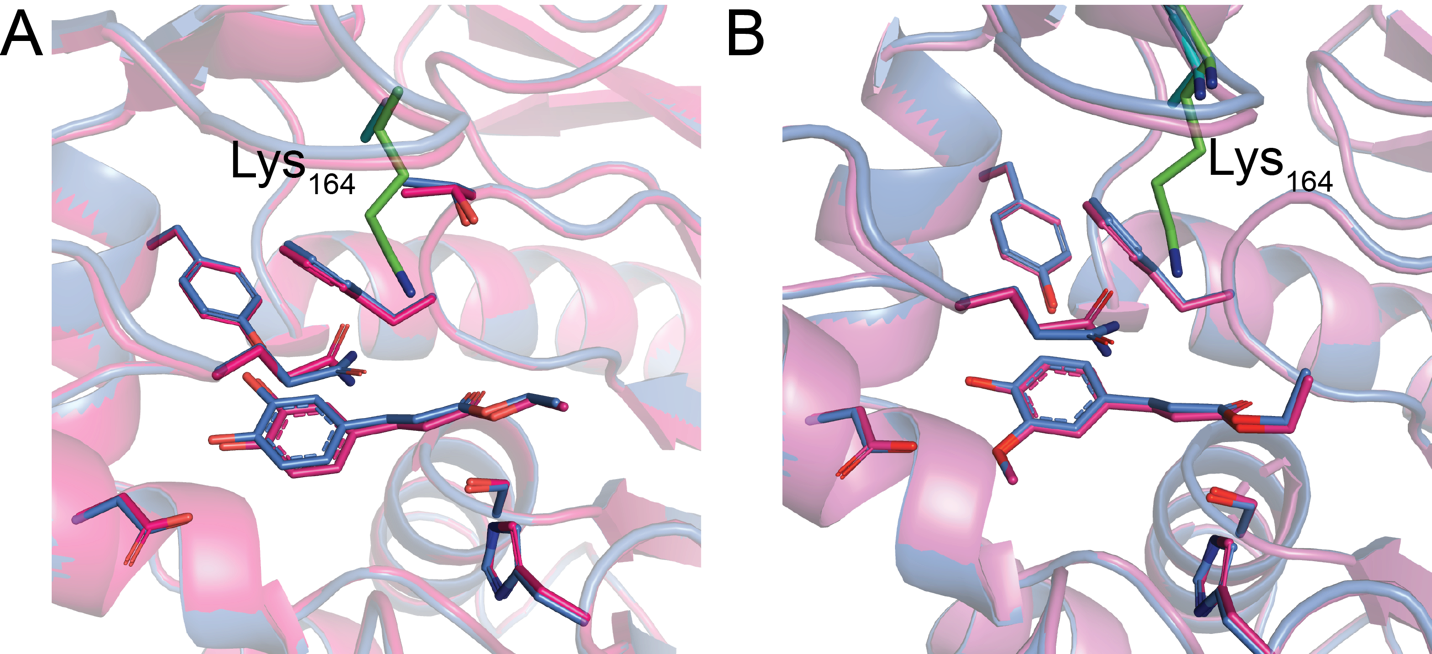
**­­**

**Figure S6.** (A) Ethyl caffeic acid and (B) ethyl ferulic acid bound to WT (blue) and K164A LhCE (pink), indicating that the substrates bind to both the wild type and mutant enzyme in the same fashion, and that Lys_164_ (green) does not interact with the respective substrates.

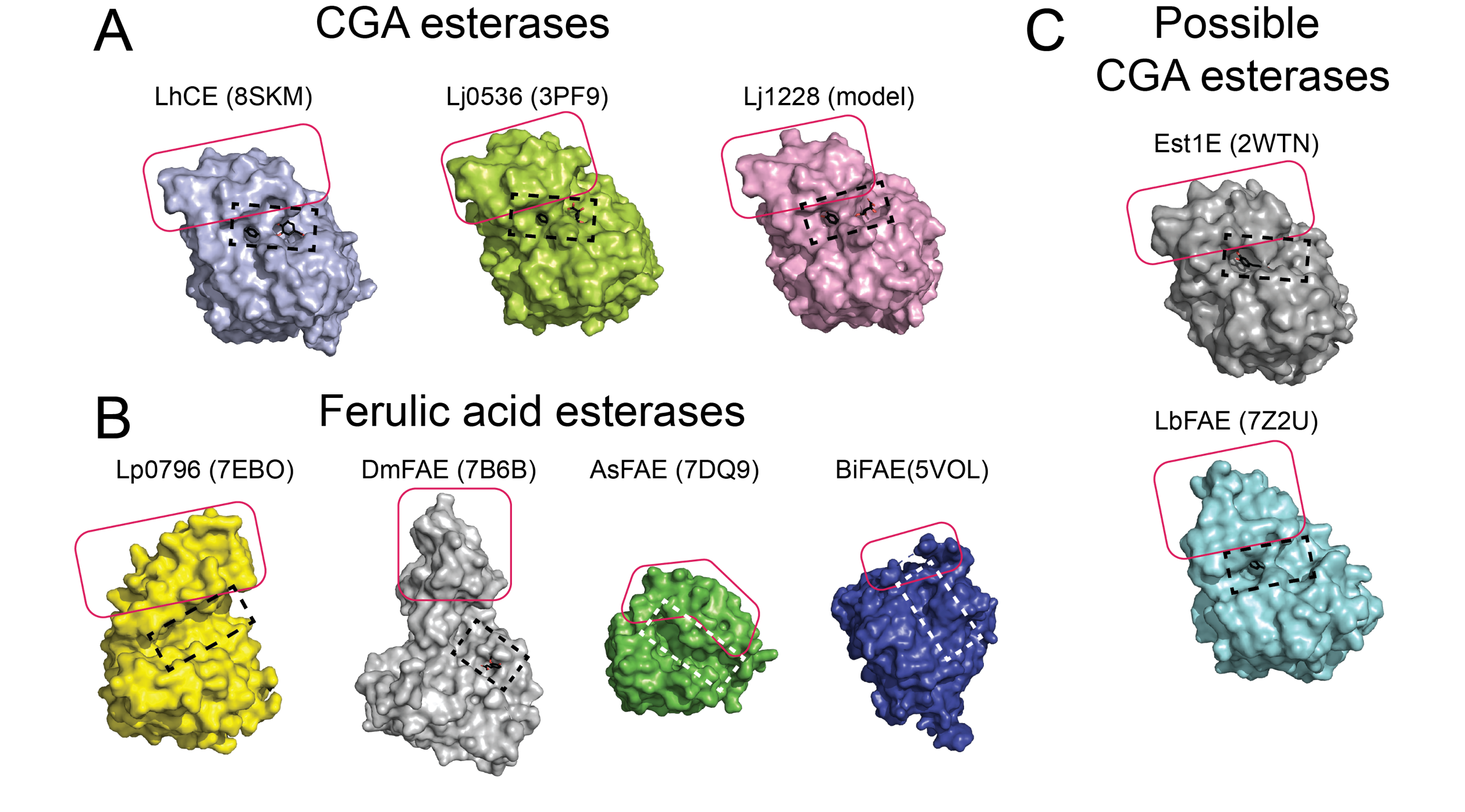

**Figure S7.** Enlarged view of main text figure 5 showing the surface representations of CGA bound to LhCE (8SKM), Lj0356 (3S2Z), and Lj1228 (Alphafold model).CGA is colored black with oxygen atoms colored red.

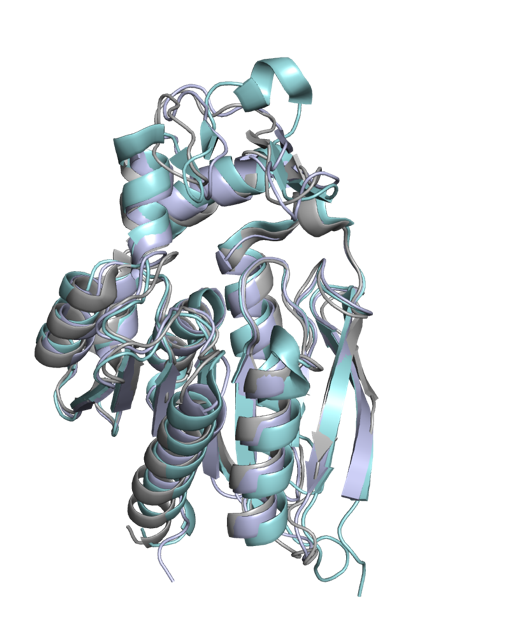

**Figure S8.** Alignment of LhCE, Est1E (2WTN) and LbFAE (7Z2U) indicating that the three proteins are very similar. The colors are: LhCE, blue; Est1E, grey; LbFAE, teal.
